# Analysing economic costs of invasive alien species with the invacost R package

**DOI:** 10.1101/2020.12.10.419432

**Authors:** Boris Leroy, Andrew M. Kramer, Anne-Charlotte Vaissière, Melina Kourantidou, Franck Courchamp, Christophe Diagne

**Affiliations:** Unité Biologie des Organismes et Ecosystèmes Aquatiques (BOREA, UMR 8067), Muséum national d’Histoire naturelle, Sorbonne Université, Université de Caen Normandie, CNRS, IRD, Université des Antilles, Paris, France; University of South Florida, Department of Integrative Biology. Tampa Fl, 33620, USA; Université Paris-Saclay, CNRS, AgroParisTech, Ecologie Systématique Evolution, 91405, Orsay, France; Institute of Marine Biological Resources and Inland Waters, Hellenic Center for Marine Research, 164 52, Athens, Greece; Department of Sociology, Environmental and Business Economics, University of Southern Denmark, 6705, Esbjerg, Denmark

**Keywords:** biological invasions, drivers of change in biodiversity, economic costs, economic impacts, ecosystem services, invasive alien species

## Abstract

1. The reported costs of invasive alien species from the global database InvaCost are heterogenous and cover different spatio-temporal scales. A standard procedure for aggregating invasive species cost estimates is necessary to ensure the repeatability and comparativeness of studies.
2. We introduce here the `invacost` R package, an open-source software designed to query and analyse the InvaCost database. We illustrate this package and its framework with cost data associated with invasive alien invertebrates.
3. First, the `invacost` package provides updates of this dynamic database directly in the analytical environment R. Second, it helps understand the heteregoneous nature of monetary cost data for invasive species, processes to harmonize the data, and the inherent biases associated with such data. Third, it readily provides complementary methods to investigate the costs of invasive species at different scales, all the while accounting for econometric statistical issues.
4. This tool will be useful for scientists working on invasive alien species, by (i) facilitating access to and use of this multi-disciplinary data resource and (ii) providing a standard procedure which will facilitate reproducibility and comparability among studies, one of the major critics of this topic until now. It should facilitate further interdisciplinary works including economists and invasion ecology researchers.

## Introduction

The economic costs of invasive alien species (IAS) are a case of heterogeneous data with different spatio-temporal scales that pose issues for global or comparative studies (Diagne, Catford, Essl, Nuñez, & Courchamp, 2020; Diagne, Leroy, et al., 2020b). Yet such studies are needed because biological invasions are a major threat to biodiversity which receive insufficient attention from decision makers and the general public (Courchamp et al., 2017). Adequately addressing the costs of biological invasions requires being able to respond to a large array of questions, such as: how are costs distributed across space, time, taxonomic groups, and economic sectors? How have these costs evolved over the last decades and can they be expected to evolve for the decades to come? How do damage and loss costs compare to management expenditures?

The absence of a standard procedure to standardize cost values for IAS may lead to the development of idiosyncratic and heterogenous methods, resulting in a lost opportunity for the repeatability and comparativeness of studies. A promising solution lies in open-source software providing frameworks to openly share data and methods altogether (e.g., Michener & Jones, 2012). Therefore, we developed the `invacost` R package as a tool to query and investigate *InvaCost* economic costs of IAS worldwide (Diagne, Leroy, et al., 2020a). This database is global in extent and covers many taxonomic groups, ecosystem types, activity sectors, and temporal and spatial scales. The `invacost` R package and its framework have already been used, thus far, in 29 publications to describe the economic costs of biological invasions at multiple spatial scales (e.g., Diagne et al., 2021; Haubrock et al., 2021).

We developed the `invacost` R package with three objectives. The first was to provide the up-to-date database directly into R, relieving users from the burdens of compatibility issues and errors associated with loading such a large dataset in R. Second, the package helps users understand the nature of monetary cost data for IAS and the inherent biases associated with such data with a step-by-step tutorial provided with the package. The third objective was to provide two complementary ways to analyse these data. One is a standard method to derive cumulative and average cost values over different periods of time, with relevant visualisation methods. The other derives the trend of costs over time with different modelling techniques accounting for the statistical issues of such econometric datasets, such as non-linearities, heteroskedasticity, temporal autocorrelation, and outliers. By meeting these objectives this software provides widespread access to these data and facilitates comparisons across studies in a straightforward manner. However, we strongly recommend working with the invacost data in interdisciplinary teams that incorporate social science expertise (i.e., economics) in order to match each specific problems with the most suitable methodological choices in handling and avoiding improper use of the data. For maximum flexibility for addressing individual researcher needs, we encourage users where necessary to duplicate the package source code and adapt it to their needs, for example altering the standardization across currencies. This possibility to adjust the code as best suited to one’s needs, coupled with the necessary economic expertise allows for flexibility and versatility in answering different questions using the necessary tools and suitable conditions.

In the following sections, we describe the rationale and methods implemented for these objectives along with relevant literature. We do so with the illustration of a simple case study on the global monetary costs caused by invasive invertebrates (i.e., all non-chordate animals).

### The invacost R package

The package requires a standard installation of R (version≥4.0.0) and is available on the Comprehensive R Archive Network (see Appendix A.1 for a code example). Upon installation eight dependencies will be automatically installed: dplyr (Wickham, François, Henry, & Müller, 2020), earth (Milborrow, Derived from mda:mars by Hastie T and Tibshirani R, & Uses Alan Miller’s Fortran utilities with Thomas Lumley’s leaps wrapper, 2019), ggplot2 (Wickham, 2016), lmtest (Zeileis & Hothorn, 2002), mgcv (Wood, Pya, & B, 2016), quantreg (Koenker, 2020), robustbase (Maechler et al., 2020), sandwich (Zeileis, 2004) and scales (Wickham & Seidel, 2020). All the package code is open-source, available on the GitHub repository, where users can also contribute or submit issues: https://github.com/Farewe/invacost. All objects created in the package have associated generic functions, meaning that if users want to see their object in the console or plot it, they will get a user-friendly output with useful information or see appropriate graphical representation of their results (Fig. 1), designed on the basis of recommended practices for data presentation, especially at small sample sizes (e.g., Weissgerber et al. 2015). In addition, output objects are lists composed of the necessary elements for reproducing the results: input data, chosen arguments, and analysis results. These objects can be stored (e.g. with `saveRDS`) and used as electronic supplementary material to ensure replicability.

**Figure 1.**
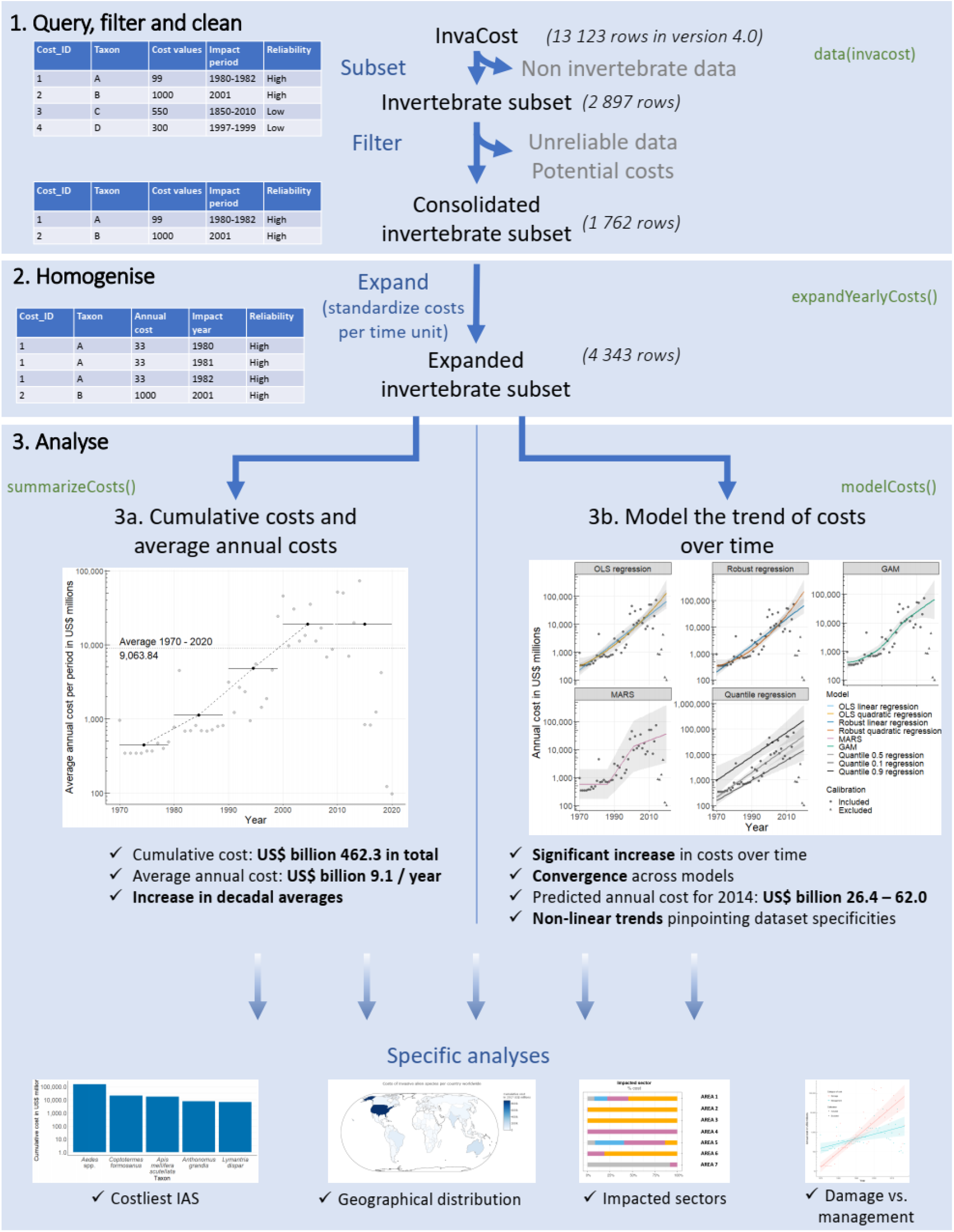
Conceptual framework of analysing monetary costs of biological invasions within the `invacost` R package. We illustrate this framework with the example of a subset of the database (invertebrates, i.e., all non-chordate animals). The framework depicts the three objectives detailed in the manuscript. We indicated in green the functions used in the `invacost` package. In 1 and 2, we illustrate with simplified tables how the structure of the database changes as the database is subsetted, filtered and then expanded. The cost over time graphs in 3a and 3b illustrate the native graphical outputs implemented in the package.

### Objective 1 – Querying, cleaning and filtering the *InvaCost* database

The *InvaCost* database is a dynamic database where existing information can be corrected and new data regularly added. Every new release of the database is checked for errors and inconsistencies with dedicated testing procedures in the package. The latest version of the *InvaCost* database (accessible at https://doi.org/10.6084/m9.figshare.12668570) is shipped with every release of the R package. The database can be accessed with the command `data(invacost)`. To make sure that users understand the database and the package, we provide a step-by-step tutorial with thorough explanations on the GitHub repository (https://github.com/Farewe/invacost).

The database loaded in R contains over 60 fields (see here for a full description https://doi.org/10.6084/m9.figshare.12668570 and here for frequently asked questions https://farewe.github.io/invacost_FAQ/, such as *(i)* how to collate data from the literature to the database, *(ii)* how double-counting is managed, or *(iii)* caveats and avenues for further improvement). These fields include, for example taxonomic information on the focal invasive taxa or geographic information on the impacted area, which enable convenient filtering within R (using ‘`subset()`’ for example) to refine the database into a subset relevant to specific research questions. To facilitate reproducibility of published analyses, we provide the function `getInvaCostVersion` to roll back the database to previous major releases (this function currently includes five releases: 1.0, 2.0, 2.1, 3.0 and 4.0).

The diversity of sources, cases and methods included in *InvaCost* will typically require users to make methodological choices about filters to apply to the database (e.g., reliable vs. unreliable sources, potential vs. observed costs) and about the costs to use (e.g., type of currency conversion factors, spatial scale of the study). We provide caveats and associated recommendations in Appendix B on these necessary choices, and everything is detailed step-by-step in the online tutorial of the invacost R package (https://github.com/Farewe/invacost).

In our example on the global costs of invasive invertebrates, we chose to filter out less reliable cost estimates and potential costs, to focus only on observed costs, which yielded a consolidated invertebrate subset of 1 762 cost estimates (Fig. 1.1, see Appendix A.2 for code).

### Objective 2 – homogenization of costs: expression in annual costs and expansion to their relevant time periods

Once the relevant filters have been applied to the database, extracting meaningful cost estimations from the resulting subset of cost records requires accounting for the fact that cost entries in the database have different temporal coverages: entries can be one-time costs, annual costs with repetitions over multiple years, or total costs of impacts which spread over multiple years. Therefore, to be comparable, cost estimates must be homogenized with a two-step process. First, they must be expressed with the same temporal unit, where the most relevant choice is annual costs. This step is already accounted for in the database with fields containing “cost_estimate_per_year” in their names. Second, once they have been homogenized on an annual basis, costs must be applied to their relevant time periods, i.e. repeated for each year over which the monetary impact was reported. This step is performed with the `expandYearlyCosts` function. This function relies on the fields indicating the starting and ending years of the annual costs. For example, reference ID 1619 reports a cumulative eradication budget of € 550,000 for *Anoplophora glabripennis* in France between 2003 and 2008. A preliminary step, already included in the *InvaCost* database standardizes the costs into a common currency. That is, conversion from local currency to US Dollars (USD) using the exchange rate or, for a better consideration of the difference of price levels among countries, the Purchasing Power Parity (PPP) and then inflation into 2017 USD (see Diagne et al. 2020 or the online tutorial of the invacost R package for details). For the purposes of the standardization Diagne et al. (2020) chose to first convert the costs from the local currency to USD and then use the appropriate inflation rate. It is important to note that the order of this process matters in determining the ultimate cost and that reversing it (i.e., inflating first in the local currency and then converting in USD) may lead to different numerical results. In the aforementioned case of ID 1619, this yields an annual cost of 2017 USD 136,437 for that period. The expansion step implemented in our package replicates this standard annual cost over each year of the impact period (2003-2008, Fig. 1.2). The costs are not expanded for the database by default because the database is easier to distribute in the compact form and because expanding the costs requires decisions which should be assessed by the user, depending on the research question addressed.

The expansion step requires adequate information with respect to the beginning and ending years of cost impacts. However, information on the beginning and ending years was not directly provided in the literature sources of monetary costs for 23% of entries in the database (2,166 rows of data). Therefore, for each source for which it was not available, educated guesses were made on the probable starting and ending years, and included these guesses in the columns “*Probable_starting_year_adjusted*” and “*Probable_ending_year_adjusted*” columns (Diagne, Leroy, et al., 2020b). Because these columns are based on conservative assumptions (e.g., the ending year of costs does not extend beyond the publication year), they should limit over-estimation; hence, it is recommended using these columns (see discussion and Extended Data Fig. 6 in Diagne et al., 2021). Consequently, this process requires removing any cost entry for which the period impact could not be extracted from the source material. Once the homogenization step has been performed on all cost entries in the user’s consolidated subset of the database, extractions and analyses can be performed to explore the patterns of costs of IAS. In our example, after expansion the data on invertebrate costs contained 4,343 rows (Fig. 1.2, code in Appendix A.3), each representing a single year of costs for a specific species/group.

### Objective 3a – Estimating the cumulative and average reported costs of invasions

The first method to analyse monetary costs of IAS consists of calculating the cumulative and average costs over time using cost estimates, as they appear in the filtered and homogenized material. These costs can be investigated on an annual basis, over the entire period covered by the database, or over a series of time intervals to account for the evolution of costs over time. All these options are performed simultaneously with the function `summarizeCosts`. First, this function calculates the sum of costs for each year of the time period requested by the user. That is by default, from 1960 (the first year with available conversion factors necessary to standardise the costs into 2017 USD as previously mentioned using exchange rates or PPP and inflation rates available through the WorldBank website for example) to the last year of the dataset. Second, it computes the cumulative total costs and average annual costs over the requested period. Last, it computes cumulative and average annual costs for user-defined time intervals (by default, 10 years) in the requested period.

A typical usage of this function is to automatically derive cumulative or average costs for specific subsets of the database, such as for specific geographical areas, type of cost, or on a per-species basis (see the detailed example on how to derive per-species cumulative cost estimates in the online tutorial https://github.com/Farewe/invacost#example-on-many-subsets-all-taxaspecies-in-the-database). In our example on the costs of invasive invertebrates, the function yielded a cumulative cost of 2017 USD 462.3 billion for the 1970-2020 time period, which corresponded to an annual cost of 2017 USD 9.1 billion per year (Fig. 1.3a, code in Appendix A4).

### Objective 3b – Modelling the trend of monetary costs of invasive alien species over time

The second analytical method implemented in the package consists of modelling the long-term trend in economic impacts of IAS by fitting models of annual costs as a function of time. Such a modelling approach is appealing because it accounts for the dynamic nature of costs, and can reliably estimate the evolution of the reported costs of IAS over time, along with estimations of uncertainty. The package implements such a modelling procedure in the `modelCosts` function, which includes four different modelling techniques with specific parameterisation resulting in a total of nine models fitted (Table 1). We chose these different statistical methods because they are complementary in their description of the trend of costs over time, and robust to the statistical issues of econometrics data: heteroskedasticity, temporal autocorrelation and outliers. We expect one or more will fulfil the general needs of most users (Table 1).

**Table 1.**
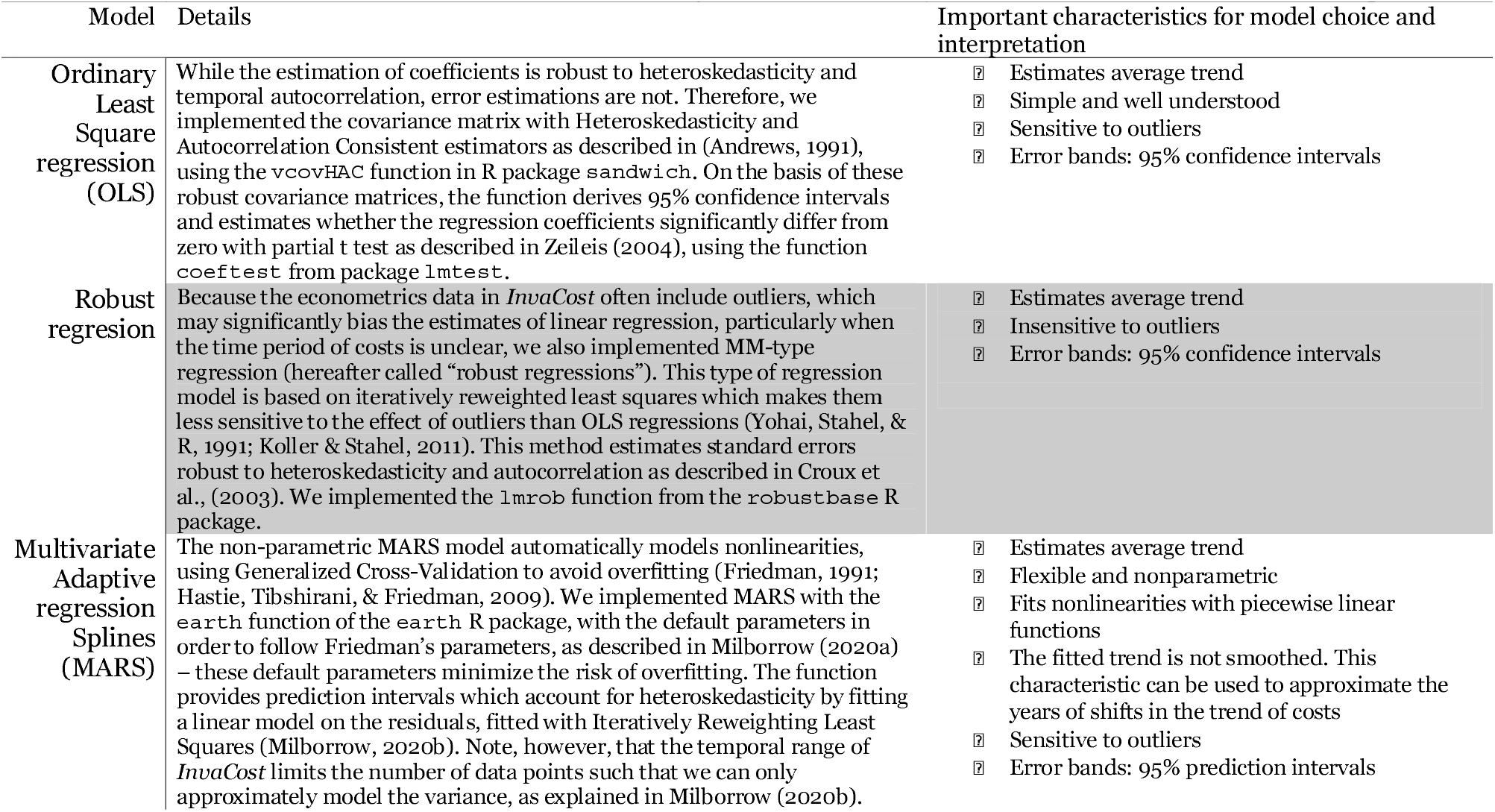
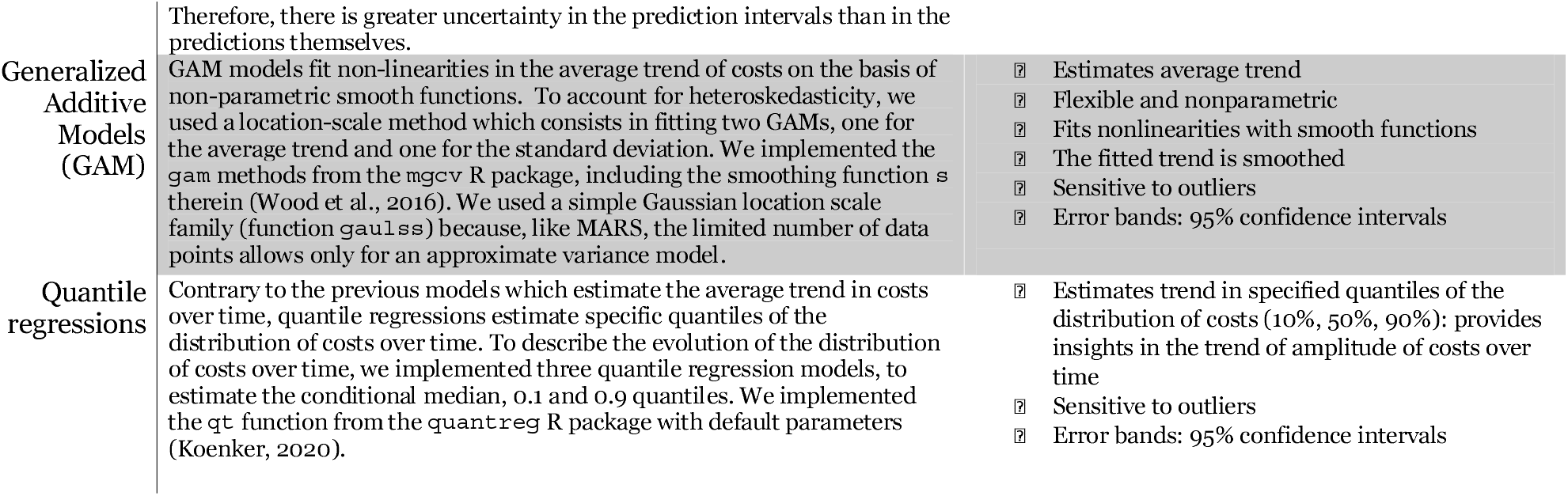
Models implemented in the `invacost` R package (function `modelCosts`), details of their implementation, and summary of their characteristics to assist in model choice and interpretation.

The fitting of these different models provides a description of the linear (with or without outlier correction) and non-linear patterns in the average trend of costs over time, as well as linear trends in the distribution of costs over time. Depending on their objectives, users can either choose one model that has characteristics fitting their question (Table 1), or compare the results of several models by analysing their convergence or divergence to describe the trend of costs over time (keeping in mind that this is affected by data characteristics). The output of the `modelCosts` function includes all the fitted models with their parameters, a table with predicted values per model over the temporal range chosen by the user, as well as diagnostic tools, such as the summary statistics specific to each model and the root mean square error between observations and predictions. The object also includes the formatted input data and parameters for reproducibility. Several parameters can be modified, including the temporal range of data to use; transformations to apply to cost values beforehand (e.g. by default, costs are log10-transformed); weights or a threshold to reduce the impact of years with incomplete data. For example, there is a lag between the occurrence of a cost and its reporting and publication in the literature. This time lag impacts the most recent years, which consequently constitute obvious outliers with the latest annual costs significantly lower than the rest of the data, a pattern pervasive to all subsets of *InvaCost*. Users can account for this incompleteness of data either by investigating results of models robust to outliers (e.g., robust regressions), by defining an optional threshold to exclude the most recent years from calibration, or by applying optional weights to reduce the influence of incomplete years on model calibration (as illustrated in examples of the online tutorial). When users are satisfied with their models and want to export results to prepare a manuscript, we provide the function `prettySummary` to export the main statistics for each model into a conveniently formatted table.

In our example on invertebrates, we excluded from model calibration all cost values from 2015 onwards, because they constituted obvious outliers with a sudden drop of two orders of magnitude (Fig. 1.3b, code in Appendix A.5). We confirmed these outliers by investigating the robust regressions which had set the weights of years above 2015 to zero (see also discussion in Supplementary Methods 1 in Diagne et al., 2021). We found significant increases in costs over time, with convergent predictions among modelling methods suggesting annual costs in 2014 of 2017 USD billion 26.4 to 62.0. Models MARS, GAM, and quadratic regressions identified non-linear patterns in costs, pinpointing an acceleration in cost increase from the mid-1980s onwards. Quantile regressions for 0.1 and 0.9 quantiles had slightly distinct slopes suggesting an increase in between-year cost amplitude over time.

## Conclusion

In conclusion, we hope that the `invacost` R package will be beneficial to researchers and stakeholders, by providing the most up-to-date version of the global database on economic impacts of IAS directly in R - with a series of standard and robust methods to extract, analyse, and compare cost data. This is the first R package to address global costs of IAS, but it follows the philosophy of similar software facilitating access to large datasets across disciplines (e.g., letsR, Vilela & Villalobos, 2015). A next step to promote access to a larger audience, beyond regular R users, will be to make all functions and the content of the package (including the database itself) accessible in a user-friendly interface, e.g., through interactive tools like Shiny apps (https://shiny.rstudio.com/). The access facilitated by the `invacost` package has already fostered new research opportunities on the impacts of IAS. For example, the package has been used in all 19 articles of the NeoBiota special issue on the geographic distribution of monetary costs of biological invasions around the world, which ultimately aim to inform decision makers at relevant scales (Zenni, Essl, García-Berthou, & McDermott, 2021).

Furthermore, the `invacost` package is an ideal tool to help achieve novel scientific assessments of invasion impacts, not merely the reported costs. Specifically, the continued improvement of the `invacost` package as well as the interoperable nature of the dynamic InvaCost database will enable: *(i)* the linkage of cost estimates to established indicators of alien impacts worldwide (e.g., GRIIS: Global Register of Introduced and Invasive Species, Pagad, Genovesi, Carnevali, Schigel, & McGeoch, 2018; SEICAT: Socio-Economic Impact Classification of Alien Species, Bacher et al., 2018) to ensure a standardised assessment of IAS impacts across regions and socio-economic sectors over time; *(ii)* investigation of the relationships between invasion costs, ecological traits, and phylogeny of IAS to better understand and predict impacts; *(iii)* movement towards predictive approaches to support, evaluate and prioritize cost-efficient management strategies according to various scenarios of invasions in our changing global environment (Lenzner et al., 2019) ; and *(iv)* opportunity to cross-analyse the costs of biological invasions with the other global drivers of change in biodiversity.

Hopefully, the `invacost` package will therefore be used as a powerful tool to complement assessments of the diversity of impacts of biological invasions, and better inform decision-makers from sub-national scales (e.g., Manfrini et al., 2021) to international assessments (e.g., the Intergovernmental Science-Policy Platform on Biodiversity & Ecosystem Services).

## Supporting information

Appendix A - Rmarkdown code

Appendix B - Additional caveats and recommendations

## Acknowledgements

We thank Anna Turbelin and Phillip J Haubrock for their assistance in proofreading and beta-testing the invacost R package. We thank Nicolas Dubos for discussions on statistics. BL, ACV and FC were funded by their salaries as French public agents. The post-doctoral contract of CD was funded by the BiodivERsA-Belmont Forum Project “Alien Scenarios” (BMBF/PT DLR 01LC1807C).

## Supplementary material

Appendix A. Code used for the analyses.

Appendix B. Additional caveats and recommendations on data filtering.

## Data availability

All data is freely available in the `invacost` R package, and the R package code is open-source and available here: https://github.com/Farewe/invacost.

## Notes

**Conflict of Interest statement** No conflict of interest.

https://github.com/Farewe/invacost

### Competing Interest Statement

The authors have declared no competing interest.

### Summary of Updates

We modified the text in several instances, in order to provide further details and guidance on the economic aspects.

