## Appendix A - Rmarkdown code for "Analysing economic costs of invasive alien species with the invacost R package"

---
title: "Appendix A. Code used for the analyses."
date: "`r Sys.Date()`"
output: word_document
---

# NOTE: bioRxiv does not allow the upload of rmarkdown (.Rmd files). 
# Please rename the file into "Appendix A.Rmd" for Rstudio to detect it as a proper Rmarkdown.

# A.1 Install and load invacost

The current stable release of invacost can be installed directly from the CRAN.
```{r eval=FALSE}
install.packages("invacost")
```

The development version of invacost can be installed with the following command
line:

```{r eval=FALSE}
install.packages("devtools")
devtools::install_github("Farewe/invacost")
```
Load the invacost package, and load the database in memory

```{r}
# Load the package
library(invacost)

# Load the database in memory
data(invacost)

# How many cost estimates in current version of invacost?
nrow(invacost)
```

Note that the manuscript was based on invacost version 4.0, so to reproduce
the results, we request this specific version:

```{r}
invacost <- getInvaCostVersion("4.0")

nrow(invacost)
```

# A.2 Querying, cleaning and filtering data for invertebrates

```{r}
library(plyr) # Install plyr if necessary
library(dplyr) 

# We can explore the descriptors with the following commands

# Kingdoms available in invacost
plyr::count(invacost$Kingdom)
# Filter invacost to the Animalia kingdom
invacost <- invacost %>%
  filter(Kingdom %in% "Animalia")

# Phyla available in invacost
plyr::count(invacost$Phylum)
# Filter invacost to invertebrates, i.e. excluding Chordata
# Note that we chose here to ignore the cost data pertaining to multiple
# phyla at the same time (e.g., "Diverse/Unspecified")
invacost <- invacost %>%
  filter(Phylum %in% c("Annelida",
                       "Arthropoda",
                       "Cnidaria",
                       "Ctenophora",
                       "Mollusca",
                       "Nematoda",
                       "Platyhelminthes"))


# Resulting number of cost estimates
nrow(invacost)
```

Now that we have invacost reduced to our list of taxa, we can start cleaning
and filtering cost data.

```{r}
# First, inspect the method reliability descriptor
plyr::count(invacost$Method_reliability)
# Filter to only keep high reliability methods
invacost <- invacost %>%
  filter(Method_reliability == "High")

# Second, inspect the implementation descriptor
plyr::count(invacost$Implementation)
# Filter to only keep "Observed" costs, omit potential costs
invacost <- invacost %>%
  filter(Implementation == "Observed")

# Next, we will inspect cost data which span multiple years but for which
# there is no starting date specified
# To do that we explore at the same time the Time range descriptor
# and the Probable starting year descriptor
uncertain.starts <- invacost %>%
  filter(Time_range == "Period" & is.na(Probable_starting_year))
# How many costs without starting date specified?
nrow(uncertain.starts)


# We do the same for uncertain endings
uncertain.ends <- invacost %>%
  filter(Time_range == "Period" & is.na(Probable_ending_year))
# How many costs without starting date specified?
nrow(uncertain.ends)


# Also, are there any costs for which it was not possible to get the time span?
# Inspect the Time range descriptor for that
unknown.periods <- invacost %>%
  filter(is.na(Time_range))
# How many costs without time span?
nrow(unknown.periods)

# If there are problematic time span costs like that, we will remove them from
# our dataset with the next lines
if(length(c(uncertain.starts$Cost_ID,
            uncertain.ends$Cost_ID,
            unknown.periods$Cost_ID)) > 0)
{
  invacost <- invacost %>%
    filter(!(Cost_ID %in% c(uncertain.starts$Cost_ID,
                            uncertain.ends$Cost_ID,
                            unknown.periods$Cost_ID)))
}

# After cleaning, we get this number of cost estimates:
nrow(invacost)

```

# A.3 Expansion step

```{r}
db.over.time <- expandYearlyCosts(invacost,
                                  startcolumn = "Probable_starting_year_adjusted",
                                  endcolumn = "Probable_ending_year_adjusted")

nrow(db.over.time)
```

# A.4 Cumulative costs

```{r}
raw.costs <- summarizeCosts(db.over.time,
                            minimum.year = 1970,
                            maximum.year = 2020)

raw.costs
```

Plot the cumulative cost estimates

```{r}
plot(raw.costs)
```
This is the basic plot as provided by default, by the invacost package.
For most users, this is sufficient.

It is possible to modify some of the parameters of this graph, such as font size
or font color, by using the regular ggplot2 commands, such as `theme()`, as
we demonstrate in the next chunk of code.


```{r}
p <- plot(raw.costs)
p <- p  +
  theme(panel.border = element_blank(),
        axis.line = element_line(color = "black"), 
        axis.text = element_text(size = 18),
        axis.title = element_text(size = 24),
        panel.grid.minor = element_blank())

```


However, advanced users may be interested in completely modifying the graphs to
fulfill their needs. In this case, we highly recommend users to 
[duplicate the source code of the package (click here to see it)](https://github.com/Farewe/invacost/blob/master/R/genericfunctions.R#L506)
and modify it to make advanced customizations to the graph. This is what
we usually do for publications, see many examples for this in the
[source code for Diagne et al. 2021, Nature](http://borisleroy.com/invacost/global_invasion_costs_scripts.html)

# A.5 Model cost trend over time


```{r fig.width = 10}
# Preliminary inspection on all data
trend.costs.exploration <- modelCosts(db.over.time,
                           minimum.year = 1970,
                           maximum.year = 2020,
                           incomplete.year.threshold = 2020,
                           final.year = 2020)
plot(trend.costs.exploration)


# Robust regression weights for the quadratic model
rrweights <- trend.costs.exploration$fitted.models$robust.quadratic$rweights
names(rrweights) <- trend.costs.exploration$fitted.models$robust.quadratic$model$Year
rrweights
```

Recent years are generally incomplete in cost data because of the delay between 
cost occurrence and reporting in the literature ([see here for a more detailed discussion](https://github.com/Farewe/invacost#correction-for-data-incompleteness-due-to-publication-la) 
and also [Supplementary Methods 1 in Diagne et al. 2021](https://static-content.springer.com/esm/art%3A10.1038%2Fs41586-021-03405-6/MediaObjects/41586_2021_3405_MOESM3_ESM.pdf)).
Therefore, it is sensible to
exclude the most incomplete years from model calibration.
The years after 2015 constitute obvious outliers visually, and the robust 
quadratic regression (which has a better R² than the linear robust 
regression) did set weights to years above 2015 to 0. Therefore, we will 
set the incomplete year threshold to 2015 in our models.

```{r fig.width = 10}
trend.costs <- modelCosts(db.over.time,
                                 minimum.year = 1970,
                                 maximum.year = 2020,
                                 incomplete.year.threshold = 2015,
                                 final.year = 2014)

trend.costs

plot(trend.costs)
```

Model summary

```{r}
summary(trend.costs)
```


Once again, we can customise the graphs, in order to make them nicer. 
In the following, we modify some of the parameters of ggplot2, to make
text more readable and adjust the legend size and position.

```{r, fig.height = 9, fig.width = 15}
plot(trend.costs) +
  theme(panel.border = element_blank(),
        axis.line = element_line(color = "black"),
        axis.text = element_text(size = 18),
        axis.title = element_text(size = 24),
        panel.grid.minor = element_blank(),
        strip.text = element_text(size = 18),
        legend.text = element_text(size = 18),
        legend.title = element_text(size = 18),
        legend.position = c(.85, .25))

```
