## Appendix B - Additional caveats and recommendations for "Analysing economic costs of invasive alien species with the invacost R package"

**Appendix B. Additional caveats and recommendations on data filtering.**

We emphasise here that users ought to inspect their filtered subset of *InvaCost* in order to grasp the nature, diversity and potential biases or limitations present	in the monetary	cost estimates provided. Indeed, as detailed in the *InvaCost* data paper (Diagne, Leroy, et al., 2020a), the database is a compilation of the data available in the literature. Such a compilation necessarily results in cost estimates that differ in design (e.g. different evaluation methods and objectives) and reporting (e.g., different spatial scales; single versus multiple species), and are incomplete, which makes any synthesis of the costs susceptible to bias (see discussion in Diagne et al., 2021). Consequently, we provide two vital recommendations to users. First, based upon their investigation of the data, users should consider applying consistency filters to their subset, such as e.g. filtering all cost estimates to the same spatial scale (as indicated in the spatial scale descriptor). Second, users should interpret the results of their analyses (such as the analyses presented in objective 3) within the limits of what they are, i.e. a snapshot of the available knowledge at the time of analysis. By that, we mean that the synthetic quantities	derived from the database reflect the inconsistent and incomplete data found in the literature; therefore, candidate hypotheses to explain the observed patterns must always include potential biases related to the underlying cost data. Ideally, considering the aforementioned and the diversity in the nature of costs reported in each study comprising InvaCost, we recommend incorporating social science expertise (i.e. economists) to appropriately address the specifics of the research question(s) that one seeks to answer.
